# Modeling the Hemodynamic Response Function for Prediction Errors in the Ventral Striatum

**DOI:** 10.1101/800136

**Authors:** Gecia Bravo-Hermsdorff, Yael Niv

**Author notes:** Corresponding author: Gecia Bravo Hermsdorff, Address: Princeton Neuroscience Institute and Psychology Department, Princeton University, Princeton, NJ 08544, USA.

## Abstract

To make sensible inferences about neural activation from fMRI data it is important to accurately model the hemodynamic response function (HRF), i.e., the hemodynamic response evoked by a punctuate neural event. HRF models have been derived for sensory areas, where it is relatively clear what events cause a neural impulse response. However, this is obviously harder to do for higher order cortices such as prefrontal areas. Therefore, one HRF model is commonly used for analyzing activity throughout the brain, despite the fact that hemodynamics are known to vary across regions. For instance, several fMRI studies use a canonical HRF to analyze ventral striatum (VS) activity where converging evidence indicates that reward prediction error signals drive neural activity. However, the VS is a target of prominent dopaminergic projections, known to modulate vasculature and affect BOLD activity, suggesting that the HRF in the VS may be especially different from those in other brain areas. To address this, we use data from an experiment focused on learning from prediction-error signals to derive a VS-specific HRF model (VS-HRF). We show that this new VS-HRF increases statistical power in model comparison. Our result is of particular relevance to studies comparing computational models of learning and/or decision making in the VS, and for connectivity analyses, where the use of an (even slightly) inaccurate HRF model can lead to erroneous conclusions. More broadly, our study highlights the importance of the choice of HRF model in determining the significance of the results obtained in classical univariate fMRI analysis.

## Introduction

The advent of fMRI revolutionized cognitive neuroscience by providing the apparatus for noninvasive spatial mapping of human cognition to brain regions. In particular, the use of behavioral computational models in conjunction with fMRI data (an approach known as model-based fMRI) is a powerful paradigm to study human learning and decision-making (O’Doherty et al., 2007, Montague et al., 2007). Several of these studies focus on analyzing ventral striatum (VS) activity, which is largely believed to reflect momentary differences between expected and obtained outcomes, i.e. reward prediction error signals. These signals are essential quantities to nearly all models of learning and choice, thus analyzing VS activity is particularly informative about the neural mechanisms underlying learning and decision-making (O’Doherty et al., 2006, Hare et al. 2008, Daw et al, 2008, Niv et al., 2012).

fMRI analysis utilizes the blood oxygenation level dependent (BOLD) signal to make inferences about the underlying unobserved neural activation. For these inferences to be accurate, it is important to correctly model the hemodynamic response function (HRF), i.e., the hemodynamic response evoked by a punctuate neural event (Boynton et al., 1996, Lindquist et al, 2009). These analyses model the BOLD response as a linear time-invariant (LTI) system; in particular, they assume that the BOLD response is proportional to the neural activity convolved with the HRF. HRF models have been mapped for sensory cortical regions, where it is relatively clear what events generate a neural impulse response (Boynton et al., 1996). However, this is harder to do for higher order cortices such as prefrontal areas, and common software packages for fMRI BOLD signal analyses typically use a single fixed HRF for the whole brain. For example, the default HRF in the SPM package peaks at 5 seconds after the punctuate event and consists of a mixture of two gamma distributions, while FSL uses a single gamma distribution that peaks at 6 seconds with 3 seconds standard deviation.

Using a “canonical” HRF for the whole brain allows modeling of neural responses in higher order cognitive regions, for which the stimuli or states that generate a neural impulse response are not yet known. However, hemodynamics vary across regions and incorrectly specified HRF models can decrease statistical power and systematically bias results (Lindquist et al, 2009, Handwerker et al., 2006). This is particular true for subcortical areas, such as the VS, due to differences in white matter density, vascular density, and other relevant physiological parameters (Logothetis & Wandell, 2004). The VS, moreover, is a target of prominent dopaminergic projections, which are known to modulate vasculature and affect BOLD activity (Knutson & Gibbs, 2006), suggesting that the HRF in the VS may be especially different from those in other brain areas.

To account for region-specific hemodynamics, some studies include the partial derivatives of the canonical HRF as extra basis functions that allow for more freedom in the shape of the HRF (Friston et al, 1998, 2007). Unfortunately, this solution adds too many additional degrees of freedom to the analysis as it does not constrain all regressors for one voxel to have the same hemodynamic response function, or all voxels in one area to have similar hemodynamics. The results of these analyses are thus harder to interpret (Lindquist et al, 2009).

These considerations underscore the need for a VS-specific HRF model. This would be particularly relevant for fMRI studies of decision-making and learning—studies that frequently rely on VS activity to distinguish between important but subtle differences in computational models of learning and choice. To obtain such an HRF model, one needs fMRI data along with a good estimate of the underlying neural activity. Fortunately, as mentioned previously, converging evidence suggests that reward prediction error is the signal driving neural activity in the VS. Thus, we used VS fMRI data from an experiment focused on learning from prediction errors signals (Niv et al, 2012) to derive a VS-specific HRF model.

## Materials and Methods

### HRF modeling

#### Modeling framework

Following the common assumption in the fMRI literature (e.g. Friston et al., 2007, O’Doherty et al, 2007), we modeled the BOLD response as a linear time-invariant (LTI) system under the General Linear Model (GLM) framework.

An LTI system is completely characterized by its impulse response function (called the HRF, *hrf*(*t*), for fMRI data), which is the temporal evolution of the system’s response to a punctuate input, or delta function activation. This means that if the impulse response function is known, the output of the system to any input *I*(*t*) is simply the convolution of the input with the impulse response, *I*(*t*) * *hrf*(*t*).

A GLM models the output, which here is the measured fMRI time-series *y* (for example, the measured activity of a single voxel, or the averaged activity of all voxels in a region of interest) as a noisy linear combination of *N* input time-series *X*, which for fMRI data analyses are the events believed to drive neural activity convolved with a given HRF. The noise is assumed to be zero-mean normally distributed noise, ε∼𝒩(0, Σ). Thus, the fMRI time-series is modeled as follows:

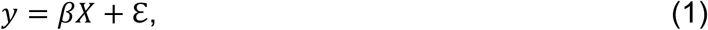

where *y* is the fMRI time-series, represented by a 1 by *T* vector, comprised of the measured discrete (*T* samples) fMRI time-series. *X* (called the design matrix) is an *N* by *T* matrix, where each row is the time-series of one of the *N* explanatory variables convolved with the HRF, the time-series of the explanatory variables are usually modeled as delta functions (for punctuate events), or boxcar functions (to represent sustained activity), and their timing corresponds to experimental events of interest, such as when a stimulus appears, when the participant makes a choice, or when a prediction error signal occurs. *β* is a 1 by *N* vector, where each coefficient controls the effect size of the corresponding explanatory variable, i.e. how much this variable contributes to the observed response. Thus, the magnitude of the coefficient *β* associated with an explanatory variable is used to determine whether that variable is significant.

Ideally, to determine the predicted activity in a region of interest, one would convolve this region’s HRF with the explanatory variables that generate a neural response in that area. However, if the HRF for a particular region is not known, it can be inferred from the data, provided that one has a good approximation for what drives the neural response in that region. For the VS, we used data from an experiment by Niv et al. (2012), which generated repeated prediction errors due to stochastic payouts and events (see detailed description below).

#### Model fitting: HRF models

We compared fits of different LTI GLMs of the VS fMRI data that used several different forms for the HRF. The functions we considered were:

1) A single gamma distribution, defined by two free parameters:

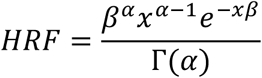

where *α* > 0 is the shape parameter, *β* > 0 the scale parameter, Γ(*α*) = (*α* − 1)!, and *x* is the temporal window of the HRF. The free parameters we fitted were the mode, *m*, and the variance, *ν*, of the HRF:

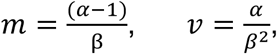

which have a one-to-one relationship with *α* and 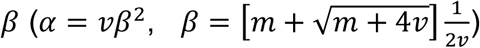 are more intuitively descriptive.

2) A weighted sum of two gamma distributions defined by five free parameters (two modes, two variances, and the relative weight or amplitude of the two components).

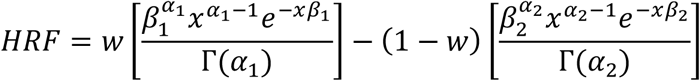

3) A finite impulse response (FIR) model, which is a nonparametric model of the HRF so as to capture any shape of response, within the limit of the frequency resolution. In the FIR model, the HRF for a given event is simply the average response over time for this event, and it has as many free parameters as the chosen resolution for the response. We used a resolution of 1 s. Although the repetition time (TR) was 2 s, we could determine the shape of the FIR with a higher resolution because the events of interest (i.e. the prediction error signals) occurred at arbitrary phases with respect to the TR onset.

#### Model fitting: Likelihood computation

The free parameters were determined by maximizing the log likelihood of the data for each model. Following other studies that estimated the HRF in other brain regions (e.g. Glover, 1999), we made the simplifying assumption that the noise is not auto-correlated. As the model of the fMRI time-course is a GLM (i.e. a linear model with Gaussian-distributed noise), the log-likelihood function ℒℒ of the model is:

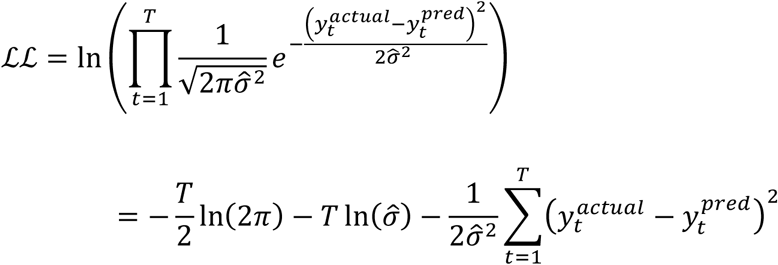

where 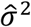 is the maximum likelihood estimator (MLE) of the variance 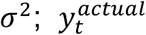 are the fMRI data sampled at time 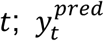 are the model predictions for the fMRI data at time *t* (according to equation 1).

For a GLM, the MLE of the variance 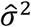 is simply the mean squared error between the data and the model (i.e. 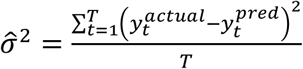), thus the log-likelihood becomes 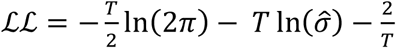. Notice that maximizing ℒℒ corresponds to minimizing the sum of the squared errors.

#### Model fitting: Optimization procedure

We fit the HRF models both to the combined data from all the participants (to find an HRF that best accounts for the population response), and to the data of each participant separately (to analyze individual variability in the VS-HRF). Due to the non-linear relationship between the HRF model parameters and the model prediction, optimization was performed numerically using the MATLAB function *fmincon*, with an “interior-point” algorithm, to minimize the negative log-likelihood function (see Nemirovski & Todd, 2008 for a review on interior-methods for optimization, and the MATLAB documentation https://www.mathworks.com/help/optim/ug/constrainednonlinear-optimization-algorithms.html for the specifics of the interior-point algorithm used). We also performed numerically unconstrained optimization using the MATLAB function *fminsearch*, which uses a Nelder-Mead simplex algorithm as described in Lagarias et al. (1998) and obtained similar results, suggesting that the values found for the HRF parameters corresponded to a global minimum as desired.

For the single gamma distribution HRF model, we constrained the mode to be between 3 to 9 s and the variance between 2 to 10 s. For the mix of two gamma distributions HRF model, we constrained the first mode to be between 2 to 10 s, the second mode to be 1 to 13 s more than the first mode, the variances to be in the range of 1.5 to 25 s, and the relative weight to be between 0 and 1. In both cases, the length of the temporal window of the HRF model did not change the results. The optimization algorithm was repeated 50 times, with initial parameter values randomly chosen within the specified bounds. For the single gamma distribution HRF model, all initializations gave nearly the same result, and agreed with those given by unconstrained optimization. Indeed, by inspecting the log-likelihood surface, we confirmed that the algorithm converged to the global maximum log-likelihood. For the mix of two gamma distributions HRF model, although it is more difficult to be sure that our results were at the global maximum as the model has more parameters, most of the fits gave the same result, including those given by unconstrained optimization.

The negative log-likelihood function’s inverse Hessian evaluated at the MLE is an (asymptotically unbiased) estimator of the asymptotic covariance matrix for the MLE fit (MacKay, 2003, Daw, 2009). Thus, we used the optimization algorithm’s numerically estimated Hessian matrix at the best fit to estimate the confidence interval for the fitted parameters. In what follows, we used 95% confidence intervals.

#### Model selection

To determine the HRF model that best fits the population, we performed leave-one-participant-out cross validation. That is, for each participant, we first fit the model parameters to the aggregate data of all other participants, and then calculated the log-likelihood of the remaining participant’s data with these parameters. We then calculated an average cross-validation score for the model as the mean log-likelihood of all the participants (when each was left out). The best model was considered to be the model with highest mean log-likelihood. In practice, cross validation can lead to overfitting (e.g. Rao et al., 2008), so we also randomly separated our data to two independent datasets: a training dataset (composed of 11 participants) that we used to fit the model parameters, and a test dataset (composed of 5 participants) that we used to evaluate the fitted models log likelihood.

To determine the HRF model that best fits an individual, we computed leave-one-session-out cross-validated log likelihoods for each participant’s data (each participant had three sessions), and we also randomly separated the participant’s data into a training dataset (composed of two sessions) and a test dataset (the remaining session).

### Experimental data

To fit a VS-specific HRF, we used VS ROI (region of interest) fMRI data from Niv, Edlund, Dayan & O’Doherty (2012). In this section we briefly describe their experiment, ROIs analysis, and data preprocessing. A more complete description of the experiment can be found in the original paper.

#### Participants

Sixteen participants (four female; age, 18–34; mean, 24 years) participated in the study. All participants gave informed consent, and the Institutional Review Board of the California Institute of Technology approved the study.

#### Experimental task

Participants made a series of choices between stimuli that paid out different amounts of monetary reward. There were five stimuli: two stimuli always paid out 0¢, one always paid out 20¢, one always paid out 40¢, and one paid out either 0¢ or 40¢ with 50% probability. Participants were not informed about these payoffs and had to learn them from trial and error. The experiment involved two types of trials that were presented intermingled: ‘choice trials’, in which participants were required to choose between two stimuli, and ‘forced trials’, in which participants were presented with only one of the five stimuli and had to choose it. The forced trials ensured that participants continued to experience the payoff associated with all of the stimuli. Choices were made immediately after the stimuli appeared on screen and reward feedback was given 5 s after the choice. There followed a variable (uniformly distributed) inter-trial interval, ranging from 2 to 5 s, after which the next trial began. If the participant took more than 1 s to respond, the trial was aborted and the inter-trial interval started. The entire experiment consisted of 234 trials (three sessions of 78 trials each), with short breaks between sessions. Before the scanning session, participants performed a training phase in which they were familiarized with the task and provided with several observations of the stimulus-reward mapping.

#### Prediction error modeling

The explanatory variable that we used in our modeling is the participant’s prediction errors during the task. In Niv et al. (2012), the data were extensively modeled and different learning models were compared against each other. We used the prediction errors obtained by the model that best fit the data, which was a risk-sensitive version of the standard temporal difference (TD) learning model (Mihatsch & Neuneier, 2002).

In the risk-sensitive TD model, the prediction error *δ* experienced at time-step *t* is given by:

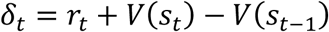

where *r*_*t*_ is the reward experienced at timestep *t* and *V*(*s*_*t*_) is the predicted value of state *s*_*t*_. Using the prediction error, the predicted value is then updated by

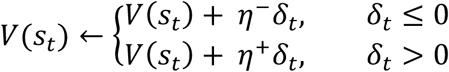

where *η*^−^ is the learning rate when the prediction error is negative, while if it is positive, the value is updated using the learning rate *η*^+^. The difference in the two learning rates translates to risk preference (*η*^+^ > *η*^−^leads to risk seeking behavior, while *η*^+^ < *η*^−^ leads to risk aversion), hence the model name. In Niv et al. (2012) the values of these parameters were fit by maximizing the log likelihood of each participant’s choice data. Here we used the best-fit parameter values from that paper to generate prediction errors for modeling the neural data.

Following Niv et al. (2012), we assumed that prediction errors occur at two time points in each trial of the experimental task: at the trial onset and at the outcome onset. At trial onset, assuming that the prior expectations on the waiting state was zero (a common assumption based on electrophysiological recordings from dopaminergic neurons; Fiorillo et al., 2003; Tobler et al. 2005), and as there is no reward at this time, the prediction error was simply the predicted value of the to-be-chosen stimulus (Niv et al., 2012). At the outcome onset, assuming that the expected reward in the trial interval state is also zero, the prediction error is the difference between the obtained outcome and the expected reward (Rescorla and Wagner, 1972).

#### ROI analysis

We used the same ROI analysis as in Niv et al (2012). The structural MRI was used to anatomically define the VS as the area bordered ventrally by the caudate nucleus, dorsally by the anterior commisure, laterally by the globus pallidus and putamen, and medially by the septum pellucidum. The border with the caudate was taken to be at the bottom of the lateral ventricle, and the border with the putamen to be at the thinnest part of the gray matter. The anterior most border was considered to be at the axial slice in which the caudate and putamen fully separated and the posterior border to be where the anterior commissure was fully attached between hemispheres.

Our analysis used the voxels in the functional space that were wholly within the VS as delineated in the higher-resolution anatomical space. We then averaged the data extracted from each of the two anatomical defined ROI (right and left VS) using singular value decomposition (SVD), as is commonly done by software packages for fMRI analysis such as SPM. This resulted in two time-course vectors of VS spatially averaged BOLD activity for each participant with samples every 2 s (i.e., every TR). From these we removed linearly modeled movement artifacts (six motion regressors: 3 translation and 3 rotation) and constant, linear and quadratic temporal trends. We then averaged the data from the left and right ROIs producing one time-series per participant.

## Results

The best-fitting VS-HRF for the population, as validated by leave-one-out cross validation and in the test dataset, was a single gamma distribution with a mode at approximately 6.0 s (standard error 0.1 s, 95% confidence interval: 5.8 - 6.2 s), and variance of approximately 3.7 s (standard error 0.6 s, confidence interval: 2.5 - 4.9 s) (Figure 1a). The best-fitting parameters for the other two models we tested (mixture of the two gamma distributions and FIR model) also had a peak at ∼6 s. Figure 1b shows a heat map of the log-likelihood surface for the single-gamma HRF, showing the peak at 6 seconds. Notably, our result is different from the canonical HRF used in SPM that peaks at 5 seconds and is comprised of a mixture of two gamma distributions, whereas we found that one gamma distribution was sufficient for modelling the VS-HRF in our dataset.

**Figure 1.**
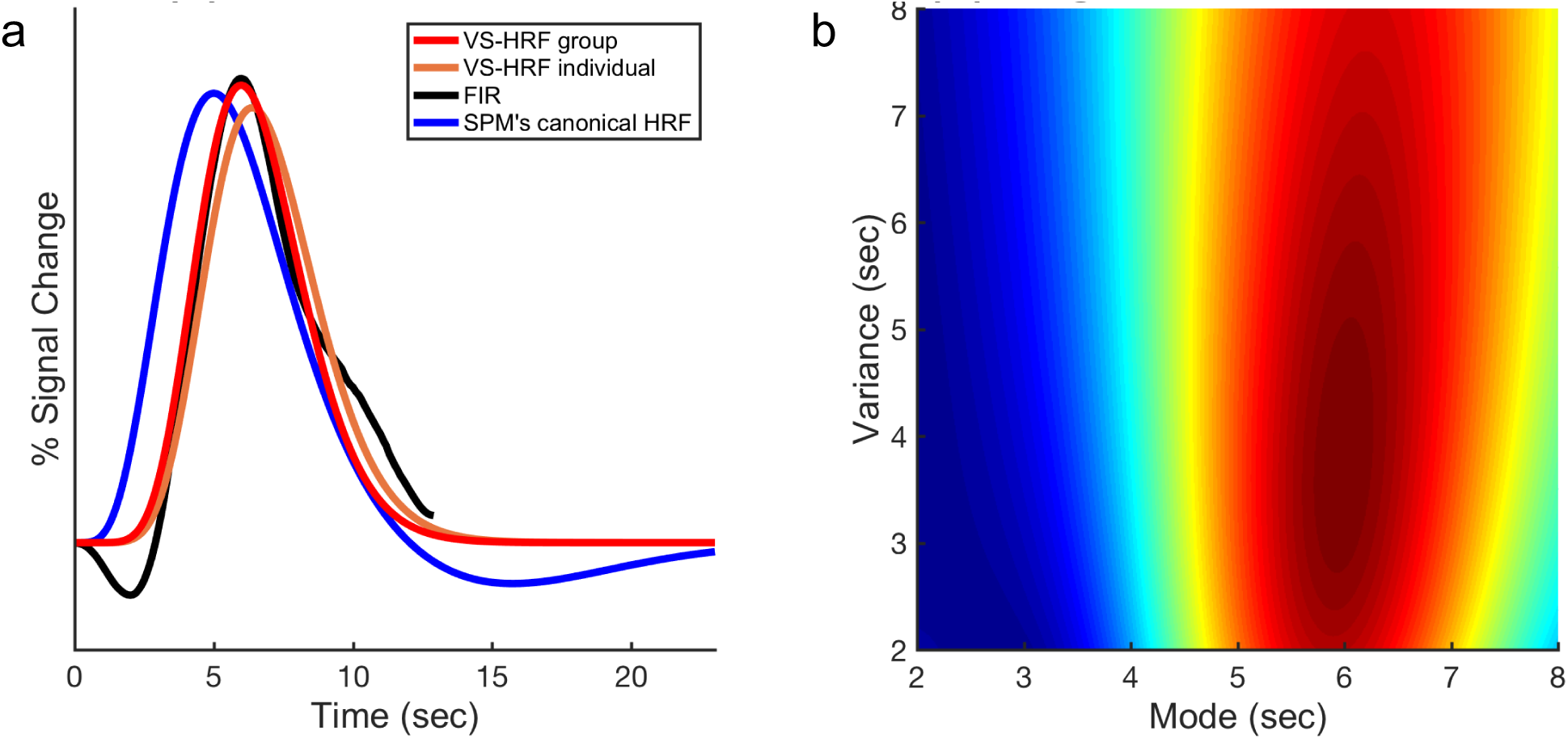
(a) VS-specific HRF. The red curve corresponds to the proposed VS-specific HRF, a single gamma distribution that peaks at 6 s and has variance 3.7 s. The orange curve corresponds to the average participant VS-specific HRF (i.e. the mean of the best fit HRF for each participant), a single gamma distribution that peaks at 6.4 s and has variance 4.1 s. The black curve corresponds to the smoothed FIR model for the population VS-HRF, which also peaks at 6 s. The blue curve, displayed for comparison, corresponds to the canonical HRF used in SPM (a gamma distribution with mode 5 s and variance 6 s minus 1/6 of a gamma distribution with mode 15 s and variance 16 s). **(b) Log**-**likelihood surface**. Heat map of the log-likelihood surface of the single-gamma HRF model calculated using the combined data from all 16 participants. Colors closer to red correspond to higher likelihood of the data.

We also fit an HRF for each participant separately. While the best-fitting VS-HRF for individual participants was also a single gamma distribution, we observed some inter-participant variability with the mean of the modes of the individual HRFs at approximately 6.4 s (standard error 0.4 s, 95% confidence interval: 5.6 - 7.2 s), and variance of approximately 4.1 s (standard error 0.75 s, 95% confidence interval: 2.6 - 5.6 s). It is conceivable that this variability in the participant-specific VS HRF was due to noise as a result of fitting the HRF to fewer data, rather than true differences between individuals.

Most fMRI studies use the general linear model (GLM) framework to determine the statistical significance of their results (Handwerker et al, 2006) by comparing the magnitude of the coefficients associated with a regressor of interest to a null (mean zero) distribution. Our results suggest that a VS-specific HRF would be better for modeling prediction errors in the VS. To measure the effect of using a VS-specific HRF in the main analysis of this dataset from Niv et al. (2012), we compared the effect size of a prediction error regressor modeling VS fMRI activity using our proposed VS-specific HRF versus using the canonical HRF from SPM. Overall, the effect size was consistently larger when using the VS-specific HRF (Figure 2), suggesting that using a VS-specific HRF significantly increases the power of statistical analysis. Interestingly, while the effect size when using an individually-fit HRF was more varied than when using the VS-specific HRF for the population, it was not significantly larger than when using SPM’s canonical HRF. This is likely due to overfitting as a result of fitting the HRF model to fewer data.

**Figure 2.**
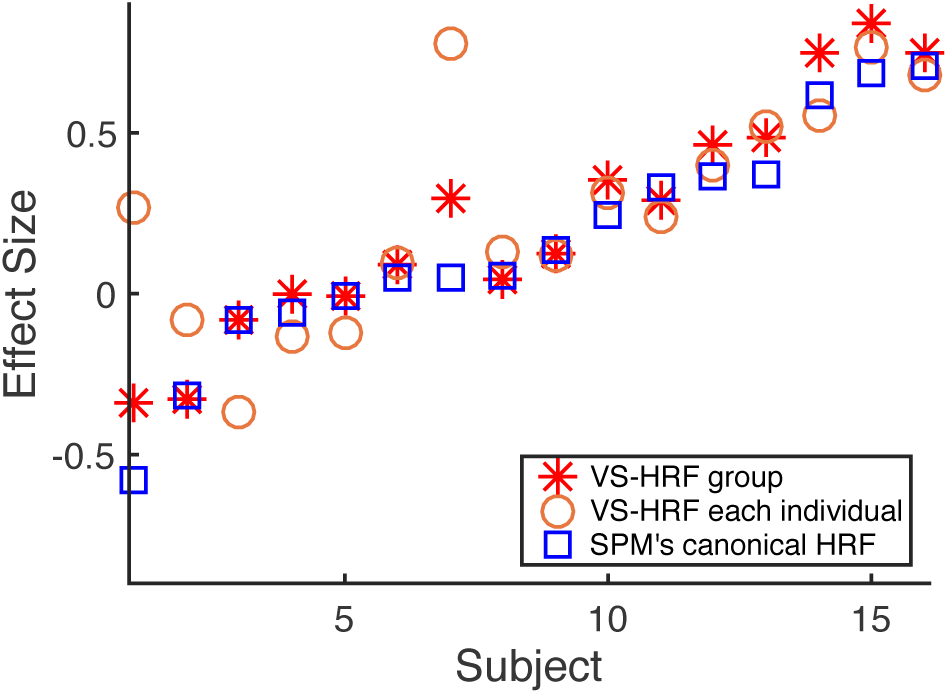
VS-specific HRF statistical power. Red asterisks denote the effect size of the prediction error regressor for each participant when the data were modeled using the VS-specific HRF; orange circles denote the effect size when using an HRF fit specifically to the participant; blue squares denote the effect size when using the SPM canonical HRF. The effect size for the model using the VS-specific was significantly larger than that of the model using SPM’s canonical HRF (p=0.003, one-tailed t-test). However, the effect size using individually-fit HRFs was more varied and not significantly better than SPM’s canonical HRF (p=0.1).

## Discussion

Incorrectly specified HRF models can decrease statistical power and systematically bias results (Lindquist et al, 2009, Handwerker et al., 2006). To remedy these problems, some studies include the partial derivatives of the canonical HRF as extra basis functions in the HRF model to allow for more freedom in the shape of the HRF (Friston et al, 1998, 2007), or use the FIR model as a flexible alternative. Unfortunately, these methods add too many additional degrees of freedom to the GLM analysis, making it harder to statistically interpret the obtained results (Lindquist et al, 2009). Thus, in this study, we leveraged the assumed correlation between prediction error signals and ventral striatal neural activity and used fMRI data from a reinforcement learning experiment focused on prediction error signals (Niv et al, 2012) to fit a VS-specific HRF.

To obtain the VS-specific HRF, we assumed that prediction errors are the effective stimuli generating an impulse function of neural response in the ventral striatum, an assumption backed by numerous studies (Hare et al, 2008; Daw et al, 2008; O’Doherty et al 2003; 2006; Van der Meer & Redish, 2011; Knutson & Gibbs, 2006). We demonstrated the superiority of the resulting VS-specific HRF in modeling data by showing that it significantly increased the statistical power of the GLM analysis in the sample dataset. This result is particularly relevant to fMRI studies of reinforcement learning and decision making that rely on detailed analysis of the VS BOLD activity to distinguish between important but subtle differences in computational models of learning and choice (O’Doherty et al, 2003; 2006; Daw et al, 2008; Hare et al, 2008; Deserno et al, 2015; FitzGerald et al, 2014). Clearly, using the wrong HRF may hinder this search.

It is also extremely important to use the correct VS-specific HRF when conducting connectivity analyses such as Granger causality and psychophysiological interactions (PPI) analysis, because inter-regional variability in the HRF that is not modeled can lead to wrong estimation of the orientation of transfer. For example, imagine that the fMRI time-series of region X is correlated with a delayed fMRI time-series of region Y. One might be tempted, as is often done in the literature, to conclude that activity in region X “drives” activity in region Y. However, if the HRF of region X is not properly modeled and the peak of its HRF is later than that of region Y, the opposite could be true. This was elegantly demonstrated by David et al., (2008), the first study to experimentally test several functional connectivity measures used on fMRI data analysis. In their study, they used a rat model of absence epilepsy, in which they knew where the spontaneous spike-and-wave discharges originated. They simultaneously performed EEG and fMRI, and subsequent intracerebral (iEEG) recordings, in strongly activated regions. Thus, they knew the actual temporal correlations of neural activity between regions, and therefore they were able to compare it with predictions from different types of functional connectivity analysis of fMRI data. They demonstrated several cases in which improper modeling of the HRF led to erroneous conclusions. Another interesting result, although not the focus of their paper, was that (in agreement with our results) they observed that the HRF was slower in the striatum as compared to other cortical regions such as V1. In theory, these data could be used to obtain region-specific HRFs for rats. However, a similarly invasive experiment in humans is not possible, and thus alternative estimates of neural activity and HRF must be used.

We used data from an fMRI experiment intended to induce specific prediction error signals. These punctate events from the model were used as a proxy for neural activity in the VS, and allowed us to derive a VS-specific HRF. As we found that this new HRF peaks later than some canonically-used HRFs, we recommend its use for future studies concentrating on the VS. More broadly, the general technique that we presented here can be applied to other brain regions. Given a putative connection between a particular experimental event and neural activity in the region of interest, a simplified experiment could be constructed to produce predictable neural events, from which an effective HRF can be derived for that region. This bespoke HRF could then be used in a more complex experiment, for which the relationship between regressor and neural activity is not known. Hopefully multiple experiments will obtain similar shapes for the effective HRF in a particular region, allowing the community to collectively generate a map of spatially varying HRFs for the whole brain. Such an endeavor will help to increase statistical power of future analyses and avoid biased results (Boynton et al., 1996, Lindquist et al, 2009), and is becoming more feasible for higher order cortical areas as their functions are slowly delineated.

## Acknowledgments

This work was supported by NIH grant R01MH098861.

## References

Boynton GM, Engel SA, Glover GH, Heeger DJ (1996). Linear Systems Analysis of Functional Magnetic Resonance Imaging in Human V1. The Journal of Neuroscience 16 (13): 4207–4221

David O, Guillemain I, Saillet S, Reyt S, Deransart C, Segebarth C, Depaulis A (2008) Identifying Neural Drivers with Functional MRI: An Electrophysiological Validation. PLOS biology 6 (12): e315

Daw N (2009). Trial by trial data analysis using computational models. Chapter in: Affect, learning, and decision making: Attention and performance xxiii

Deserno, L., Huys, Q. J. M., Boehme, R., Buchert, R., Heinze, H. J., Grace, A. A., Dolan, R., Heinz, A., & Schlagenhauf, F. (2015). Ventral striatal dopamine reflects behavioral and neural signatures of model-based control during sequential decision making. PNAS 112 (5):1595–1600

FitzGerald THB, Schwartenbeck P, Dolan RJ (2014). Reward-related activity in ventral striatum is action contingent and modulated by behavior relevance The Journal of Neuroscience 34 (4): 1271–1279

Fiorillo CD, Tobler PN, Schultz W (2003). Discrete coding of reward probability and uncertainty in dopamine neurons. Science 299: 1898–1902

Friston, K. J., Ashburner, J., Kiebel, S. J., Nichols, T. E., and Penny, W. D. (eds.). (2007). Statistical Parametric Mapping: The Analysis of Functional Brain Images. Amsterdam: Academic Press.

Friston KJ, Fletcher P, Josephs O, Holmes A, Rugg MD, Turner R. (1998). Event-related fMRI: characterizing differential responses. Neuroimage 71(1): 30–40

Glover HG (1999). Deconvolution of impulse response in event-related BOLD fMRI. Neuroimage 9: 416–429

Handwerker DA, Ollinger JM, Friston K, D’Esposito M (2004). Variation of BOLD hemodynamic responses across participants and brain regions and their effects on statistical analyses. NeuroImage 21 (4): 1639–51

Hare TA, O’Doherty J, Camerer CF, Schultz W, Rangel A. (2008). Dissociating the role of the orbitofrontal cortex and the striatum in the computation of goal values and prediction errors. Journal of Neuroscience 69: 1204–1215

Henson R, & Friston K (2007). Convolution Models for fMRI. In: Statistical Parametric Mapping: The Analysis of Functional Brain Images, pp. 178–192.

Knutson B & Gibbs EBS (2006). Linking nucleus accumbens dopamine and blood oxygenation. Psychopharmacology 191: 813–822

Lagarias JC, Reeds JA, Wright MH, Wright PE (1998). Convergence Properties of the Nelder-Mead Simplex Method in Low Dimensions. SIAM Journal of Optimization 9 (1):112–147

Lindquist DA, Meng Loh J, Atlas LY, TWager TD (2009). Modeling the Hemodynamic Response Function in fMRI: Efficiency, Bias and Mis-modeling. NeuroImage 45 (1 Supp): S187–98

Logothetis NK, Wandell BA (2004). Interpreting the BOLD signal. Annual Reviews Physiology 66:735–49

MacKay D (2003). Information theory, inference, and learning algorithms. Cambridge University Press

Mihatsch O & Neuneier R (2002). Risk-Sensitive Reinforcement Learning. Machine Learning 49(2–3):267–290

Montague, PR, King-Casas, B, Cohen, JD (2007). Imaging valuation models in human choice Annual Review of Neuroscience 29:417–448

Niv Y, Edlund J, Dayan P, O’Doherty JP (2012). Neural Prediction Errors Reveal a Risk-Sensitive Reinforcement-Learning Process in the Human Brain. The Journal of Neuroscience 32 (2): 551–562

Nemirovski SA & Todd, MJ (2008). Interior-point methods for optimization. Acta Numerica 191–234

Niv Y & Schoenbaum G (2008). Dialogues on prediction errors. Trends in Cognitive Sciences 12 (7): 265–272

O’Doherty JP, Buchanan TW, Seymour B, Dolan RJ. (2006). Predictive neural coding of reward preference involves dissociable responses in human ventral midbrain and ventral striatum. Neuron 49:157–166

O’Doherty JP, Dayan P, Friston K, Critchley H, Dolan R (2003). Temporal difference models and reward related learning in the human brain. Neuron 38 (2): 329–337

O’Doherty JP, Hampton A, Kim H (2007) Model-based fMRI and its application to reward earning and decision making. Ann N Y Acad Sci 1104: 35–53.

Padoa-Schioppa C & Assad JA (2006). Neurons in orbitofrontal cortex encode economic value. Nature 441: 223–226

Rao BR, Fung G, Rosales R (2008). On the dangers of cross-validation. An experimental evaluation. In Proc. of the 2008 Society for Industrial and Applied Mathematics (SIAM) International Conference on Data Mining

Rescorla RA, Wagner AR (1972). A theory of Pavlovian conditioning: variations in the effectiveness of reinforcement and nonreinforcement. In: Classical conditioning II: current research and theory (Black AH, Prokasy WF, eds), pp. 64–99. New York: Appleton-Century-Crofts.

Tobler PN, Fiorillo CD, Schultz W (2005). Adaptive coding of reward value by dopaminergic neurons. Science 307:1642–1645

Van der Meer M AA & Redish DA (2011). Ventral striatum: a critical look at models of learning and evaluation. Current Opinion in Neurobiology 21:387–392

Wilson RC, Takahashi YK, G Schoenbaum G & Niv Y (2014). Orbitofrontal cortex encodes a cognitive map of task space. Neuron 81(2): 267–279

